# Identification of lactic acid bacteria with reduced inflammatory and enhanced protective properties against influenza virus

**DOI:** 10.1101/2022.08.14.503913

**Authors:** Atsushi Sugimoto, Tomoe Nakamura, Ryota Chihama, Yuto Takenaka, Yuuki Sato

## Abstract

Lactic acid bacteria (LAB) exert beneficial effects on health by regulating innate immunity in the intestinal tract. Many LAB that demonstrate anti-viral properties depend on the host’s inflammatory responses; however, a few LAB strains have low inflammatory and anti-viral properties. In this study, the LAB strain D279 (NITE_BP-03645, *Latilactobacillus sakei*) was isolated from among 741 LAB strains that were analyzed for their ability to induce interleukin 12 (IL-12), and was subsequently characterized. D279 induced the highest levels of IL-12 among the screened LAB. Further, D279 significantly activated anti-viral genes and preferentially induced IFNλ *in vitro*, which plays a critical role in the epithelial tissue, conferring strong anti-influenza potency without inflammation, whereas IFNα levels were relatively lower. Administration of pasteurized D279 to mice resulted in strong anti-influenza potency, including higher natural killer (NK) cell activity, and a reduced number of viruses in the lung compared to the control. Importantly, no D279-fed mice were sacrificed during the viral infection tests. Taken together, these results suggest that D279 administration confers beneficial effects with reduced inflammation, regulates innate immunity, and may be relevant for commercial use in the future.

## Introduction

Lactic acid bacteria (LAB) have attracted considerable attention owing to growing health awareness regarding their beneficial effects. LAB act as intestinal regulators and have immunomodulatory and defense functions in the intestinal tract of the host [1]. Several studies to elucidate the defense mechanisms of various LAB strains against viruses and pathogenic bacteria have been performed to date [2]. LAB with anti-viral potency are in demand as a countermeasure against influenza infection or coronaviruses, and therefore, the identification and characterization of biogenic LAB strains is critical to further promote human health and the food industry [3].

Immunomodulation of LAB is mediated by innate immunity [4–6]. In the intestinal tract, LAB are show uptake by immune cells such as plasma dendritic cells (pDCs) or macrophages, or through microfold cells on the Peyer’s patches, leading to activation of innate immunity [7]. pDCs and macrophages recognize microorganism-associated molecular patterns (MAMP) by utilizing pattern recognition receptors (PRRs) which localize to the immune cell surface and intracellular vesicles [8–10]. Toll-like receptors (TLRs) are the most well-characterized PRRs. To date, 13 types of TLRs have been identified in humans, each of which recognizes different MAMPs [11], indicating the versatility of the immune system. TLR1, 2, 4, 5, and 6 are expressed on the plasma membrane and recognize the membranes and lipoproteins of pathogens [11]. TLR3, 7, 8, and 9 are localized to the endosomes and recognize exogenous nucleic acids such as non-methylated CpG-oligo DNA (CpG-ODN) and RNA molecules from pathogens [11].

TLRs promote the production of interleukins (ILs) and interferons (IFNs), resulting in the expression of anti-viral factors that interfere with viral replication [12]. IL-12, a heterodimeric cytokine, is produced by macrophages or phagocytic cells and plays a primary role in driving innate immunity by triggering activated NK cells [13]. Additionally, innate immunity is orchestrated by three types of IFNs: Type I (IFNα, β, and others), Type II (IFNγ), and Type III (IFNλ). Among these, IFNα and IFNλ are particularly effective at protecting against viral infection [14, 15]. IFNα is receptive to cells throughout the body and immediately induces the expression of anti-viral factors and inflammation [16]. In contrast, IFNλ, a recently discovered non-inflammatory IFN, mainly regulates epithelial cells that are at the front line of infection, in which the response is characterized by prolonged and strong induction of mucosal anti-viral factors [15, 17]. Although there is overlap between the anti-viral genes induced by IFNα and IFNλ, reports indicate that IFNλ is more effective than IFNα in protecting cells from respiratory viruses [17, 18].

Several LAB are commercially available to counteract viral infection [19–24]. Most LAB that shows anti-viral function act via the induction of inflammatory cytokines, frequently utilizing IFNα in TLR9-mediated pathways [25]. To our knowledge, one LAB alone induces immunomodulatory functions without IFNα [26]. Inflammatory cytokines can effectively suppress viral replication; however, this suppression is not always beneficial [27]. Excessive inflammation induces a cytokine storm that adversely affects recovery after influenza infection [28]. If more LAB can strongly activate innate immunity with a reduced inflammatory response, these strains can be marketed as a food material that can counteract infectious diseases. In this study, we screened LAB to identify strains with anti-influenza activity. IL-12 is an early response cytokine that acts against infection and activates NK cells to eliminate pathogens and infected cells that cause inflammation [29]. IL-12 and NK cells are effective against influenza [30, 31]. Because of these properties, IL-12 is regarded as an inflammatory and anti-inflammatory cytokine [32].

In this study, we aimed to screen LAB based on their IL-12 production capability, identify and characterize strains that induced the highest levels of IL-12, and determine whether the strain could activate innate immunity.

## Materials and Methods

### Mice

7.5-week-old male BALB/c mice were purchased from Charles River Laboratories Japan, Inc., and used for IL-12 quantification. Four-week-old female BALB/c Cr Slc mice from Japan SLC, Inc. were used for viral infection experiments because of the extensive information available on infection protocols and phenotyping methods [33]. For IL-12 quantification and influenza tests, 3 male BALB/c and 75 female BALB/c Cr Slc mice were used, respectively. Their body weights ranged from 13.2 g to 16.3 g. The detailed experimental methods and breeding conditions are described below.

### Breeding conditions

Female BALB/c Cr Slc mice were pre-bred for seven days. Weights were measured on an electronic balance on the day after obtaining mice and the day the pre-breeding period was completed. The general growth conditions of the mice were observed daily, and those that showed no significant changes in body weight or general conditions were used for grouping. Mice were bred in a room maintained at 19 to 24 °C, at 43% to 66% humidity, with 12 h of light and dark conditions (light conditions: 0600 h to 1800 h, dark conditions: 1800 h to 0600 h) at the Japan Bio Research Center. During the pre-breeding period and after grouping, animals were maintained in sterile plastic cages (W:175 × D:245 × H:125 mm) with floor mats (PaperClean Bedding, Japan SLC Inc.). The same conditions were used during the pre-breeding period and after grouping. To ensure an optimal environment, each cage was filled with environmental enrichment medium (nesting sheets, K3510, Animec, Japan). During the pre-breeding period and after grouping, mice were bred individually. Feeders were changed during feeding, and the cages, water bottles, and environmental enrichment were changed at least twice per week. The rooms were cleaned and disinfected daily. During the pre-breeding period, the animals were fed powdered feed (MF; ORIENTAL YEAST Co., Ltd., Japan) *ad libitum* within 9 months of manufacture. After grouping, the animals were fed MF or MF mixed with 5% (w/w) of the test substance. Contaminants, bacterial counts, and nutrient content in the feed were evaluated by Eurofins Food Testing Japan K.K. and ORIENTAL YEAST Co., Ltd. The drinking (tap) water was available *ad libitum*. The number of contaminants and bacteria in the drinking water were analyzed by Tohzai Chemical Industry Co., Ltd.

### Mouse groups

The mouse groups were stratified by body weight using a computer program (IBUKI, Japan Bioresearch Center Co., Ltd., Gifu, Japan), and the animals were randomly selected on the last day of pre-breeding. The mean body weights and variances in each group were approximately equal. The remaining animals were euthanized by releasing blood from the abdominal aorta under anesthesia with isoflurane on the grouping day.

### Virus preparation

Influenza virus PR8 (A/PR/8/34 H1N1) and virus-containing media were prepared by the Japan Bioresearch Center Co., Ltd. (Gifu, Japan). Briefly, the cryopreserved virus stock solution was thawed, diluted 104-fold in DMEM (D5303, Sigma-Aldrich, USA), and added to a culture flask containing MDCK cells (kindly provided by the Aichi Medical University School of Medicine Microbiology and Immunology Seminars.). To enable the virus to adsorb onto cells, the culture flask was incubated for 1 h at 37 °C in a CO_2_ incubator (MCO-170AICUV-PJ; Panasonic Healthcare, Japan).

After the virus solution was removed, DMEM with 0.25% trypsin (25300054, Thermo Fischer Scientific, USA) was added, and the cells were kept in a CO_2_ incubator until a cytopathic effect (CPE) was observed. The supernatant was centrifuged (4 °C, 780 × *g*) in an AX-310 centrifuge (TOMY SEIKO, Japan) to remove foreign substances. The viral stock solution was frozen in an ultra-low temperature freezer until use.

### Influenza infection

The influenza virus PR8 (A/PR/8/34 H1N1) suspension at a density of 2×10^3^PFU/ml was thawed, and 0.05 ml of virus-containing medium was dropped onto the nasal cavity of the mice (1.0×10^2^ PFU/mouse). The inoculation date was set as day 1. The animal experiments were approved by the Japan Bioresearch Center Co., Ltd. (Test No. 410057). Optimal measures were taken to minimize the pain and stress of the animals according to the humanitarian endpoints for the animal. All animal tests were performed and analyzed by the Japan Bioresearch Center Co., Ltd. The study was approved by the Committee of Animal Care of Japan Bioresearch Center Co., Ltd. (Test No. 410057).

### Mouse phenotyping

Mice were observed daily, and the clinical outcomes were scored once a day and are described in Table 1. The net body weight was measured once a week before viral infection. After the viral infection, body weight measurements were performed daily. Food consumption was measured twice a week. The amount of food per feeder was measured using an electronic balance on the feeding day, and the remaining amount was measured on the next feeding day. To calculate the survival rate, the emaciation state was evaluated daily according to the humanitarian endpoint, and the mice were euthanized using isoflurane anesthesia as required.

**Table 1.**
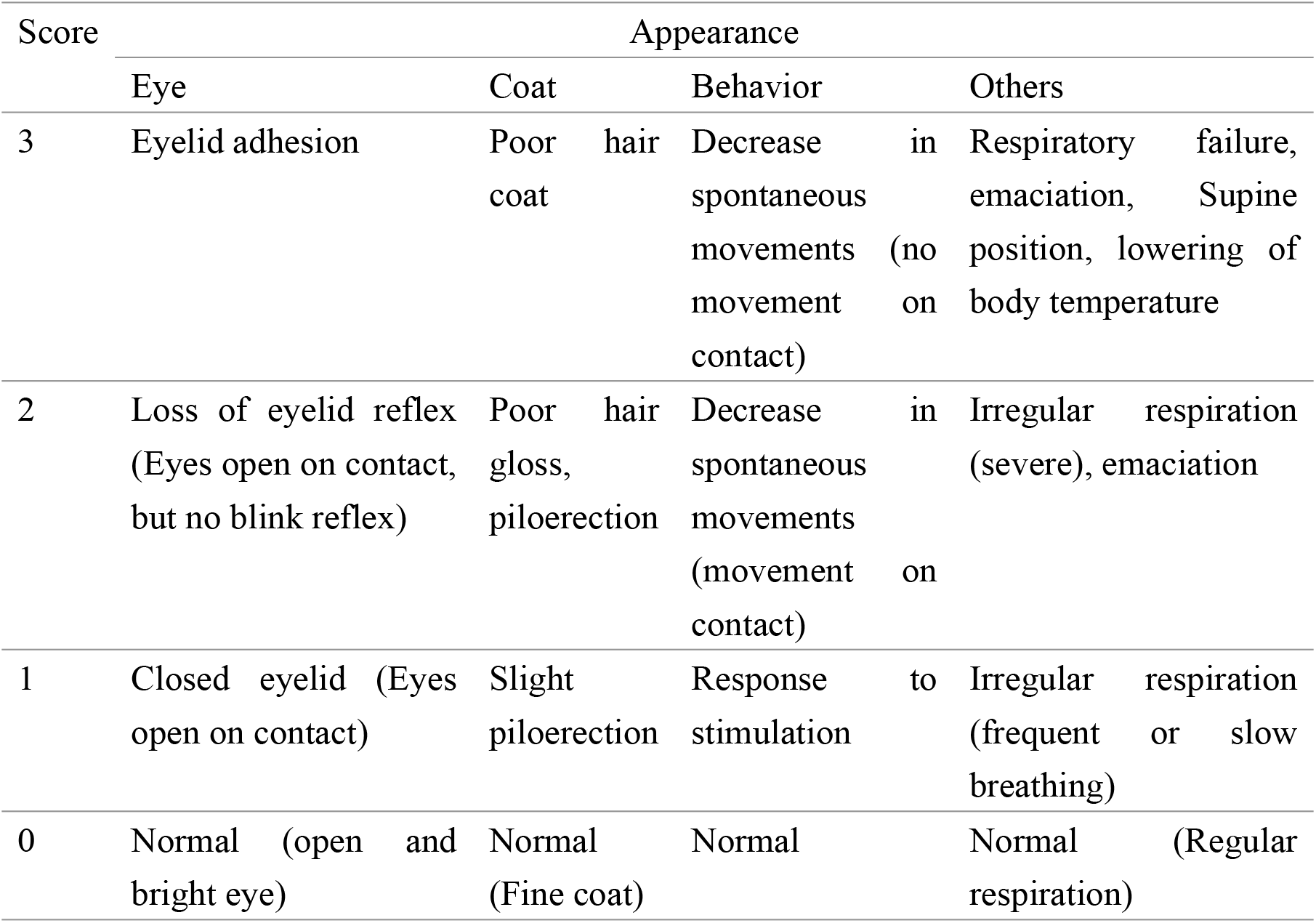
Definition of clinical scores. Clinical scores were decided based on the definitions provided in Table 1.

### IgA quantification

On day 5, mice were anesthetized and blood was collected using a syringe. Serum was obtained by centrifugation of the blood at 2000×*g* for 15 min. IgA levels were quantified using a mouse serum IgA ELISA Kit (ab157717, Abcam, UK) according to the manufacturer’s protocol.

### NK activity

NK cell activity was quantified using the LDH Cytotoxicity Detection Kit (MK401, TaKaRa, Japan) according to the manufacturer’s protocol. Briefly, the mouse spleen was excised under anesthesia on day 5 and squashed to obtain single cells in HBSS buffer (14025092, Thermo Fisher Scientific, USA). Erythrocytes were lysed by adding 2 ml of lysing solution (Ohtsuka, Japan). Mouse spleen cell cultures were diluted to a density of 4×10^6^ cells/ml and co-incubated with FITC-labeled YAC-1 cells (kindly provided by the Aichi Medical University School of Medicine Microbiology and Immunology Seminars) prepared at a density of 1×10^5^ cells/ml.

### Lung viral counts

The number of viruses in the lungs was calculated using a plaque assay. Briefly, on day 5, the mouse lungs were excised, sliced into small pieces, and homogenized in 2 ml HBSS buffer (14025092, Thermo Fisher Scientific, USA) using a Physcotron homogenizer (Microtec Co., Ltd., Japan). Homogenates were filtered using a cell strainer (pore size 100 μm, Corning) and centrifuged at 190 *×g* for 5 min to obtain the virus-containing supernatant. MDCK cells were incubated for 1 h with 0.1 ml of the virus supernatant to enable the influenza virus particles to adsorb to the cell surface. Next, a MEM-based medium containing 0.8% (w/v) agarose was overlaid onto the MDCK cells which were grown for another 2 days. The cell sample was stained overnight with neutral red (140-00932, FUJIFILM Wako, Japan) to observe the plaques and calculate the viral number. Bronchoalveolar lavage fluid was not used in the study.

### Cell culture

HT-29 cells were obtained from the European Collection of Authenticated Cell Cultures, UK (ECACC). HT-29 cells were grown with 5% CO_2_ at 37 °C in McCoy’s 5A medium (SH30270.01, Cytiva, USA) supplemented with 10% fetal bovine serum (S0250, BioWest, France) and 1% (v/v) penicillin and streptomycin (15140-122, Gibco, USA).

### LAB strains and growth conditions

A total of 741 LAB strains were selected from a collection stored at –80 °C. Each LAB was grown in MRS medium (63-6530-37, BD Difco, USA) at 30°C for 18 h. For IL-12 quantification and animal testing, LAB were grown and centrifuged at 5000 rpm for 5 min. The bacterial pellet was washed with water up to three times and sterilized at 80 C for 30 min. The LAB suspension was lyophilized (DF-05H-S, ULVAC, Japan) at –40 °C overnight to obtain a fine powder. The LAB powder was used for IL-12 quantification at a final concentration of 20 μg/ml, or mixed in the mouse feed at 0.5% (w/w).

### DNA extraction

Genomic DNA from the LAB strains was extracted using a NucleoBond AXG column and Buffer Set III (U0544A & U0603A, MACHEREY-NAGEL, Germany) according to the manufacturer’s protocol. The DNA concentration was measured using a SimpliNano spectrophotometer (29061712, Biochrom, UK) and used for lipofection.

### Enzyme-linked immunosorbent assay (ELISA)

To quantify IL-12, the spleens of the male BALB/c mice were excised and gently squeezed to obtain cells for spleen cell culture. Cells were treated with 0.75% (w/v) ammonium chloride in 17 mM Tris-HCl (pH7.7) to lyse the erythrocytes. After washing with 0.1% (w/v) BSA in PBS, cells were resuspended in RPMI 1640 medium (SH30027.01, Gibco, USA) supplemented with 10% (v/v) bovine serum (S0250, BioWest, France), and 1% (v/v) penicillin and streptomycin (15140-122, Gibco, USA). The cells were cultured with 50 μM 2-mercaptoethanol in a 96-well plate at a density of 5×10^5^ cells/well, and co-incubated with lyophilized LAB (20 μg/ml). After 48 h, the cellular supernatant was collected, and IL-12 quantification was performed. The supernatant was added to an immune plate (430341, Thermo Fischer Scientific, USA) coated with a monoclonal IL-12-B antibody (51004-R020, Sino Biological, China), and the plate was incubated at 4 °C for 18 h, followed by blocking with 5% (w/v) skimmed milk containing 0.05% (w/v) Tween-20 at 37 °C for 2 h. The plate was washed with PBS and 50 μl of the supernatant was added, and the reaction was incubated at 37 °C for 2 h. Next, 100 μl of biotinylated anti-IL-12 p35 (3456-6-250, MabtechAB, Sweden) was added and the plate was incubated at 37 °C for 2 h. The plate was washed twice with PBS, and 100 μl of streptavidin-HRP (RPN1231-2ML, Cytiva, USA) was added and the reaction was incubated at room temperature for 1 h. After washing with PBS, 100 μl of TMB substrate (5120-0047, SeraCare, USA) was added to the reaction and the plate was incubated at room temperature for 1 h in the dark. After adding the TMB stop solution (5150-0020, SeraCare, USA), the absorbance was evaluated at 450 nm using the microplate reader ARVO X3 (PerkinElmer, USA). IL-12 p70 (200-12, PeproTech, USA) was used at appropriate dilutions for the standard.

For IFNα and IFNλ quantification, HT-29 cells were cultured in 96 well plates at a density of 4×10^5^ cells/well for 24 h. Next, genomic DNA from LAB or Poly (I: C) (4287/10, R&D Systems, USA) was transfected into the cells at a final concentration of 5 μg/ml and 1 μg/ml using Lipofectamine 2000 (11668027, Thermo Fischer Scientific, USA), respectively, according to the manufacturer’s protocol. Cells were further cultured for 24 h, and the supernatant was used to quantify IFN using the VeriKine-HS Interferon α All Subtype TCM

ELISA Kit (41135-1, PBL Assay Science, USA) and the Human IL-29/IL-28B (IFNλ1/3) DuoSet ELISA (DY1589, R&D Systems, USA) kit according to the manufacturer’s protocol.

### Electron microscopy

Using electron microscopy, the phenotype of D279 cells was evaluated for whether the cells were round or rod-shaped. D279 cells were grown in MRS medium (63-6530-37, BD Difco, USA) overnight and treated with the ionic liquid HILEM IL1000 (Hitachi High-Tech, Japan) at a ratio of 1:1 for 1 h at room temperature. The mixture was dropped onto filter paper and dried until the water evaporated. After fixing the filter paper on the SEM specimen table, the sample was examined at 0.5 kV using the FE-SEM SU9000 (Hitachi High-Tech, Japan).

### Phylogenetic analysis

As D279 was not classified, the 16S rDNA sequence of D279 was analyzed using the ABI PRISM 3500xL Genetic Analyzer System (Thermo Fischer Scientific, USA), and a phylogenetic tree was created using the maximum-likelihood or neighbor-joining methods.

### qRT-PCR

Total RNA was extracted from HT-29 cells using the CellAmp Direct RNA Prep Kit for RT-PCR (3732; TaKaRa, Japan) after the introduction of genomic DNA or Poly (I: C). qRT-PCR was performed using the One Step TB Green PrimeScript PLUS RT-PCR Kit (RR096A, TaKaRa, Japan) and a Thermal Cycler Dice Real Time System TP800 (TaKaRa, Japan). *GAPDH* was used as an internal control. Primers were synthesized by Integrated DNA Technologies (USA). The specific primer sequences were as follows: *GAPDH* (Forward: 5′-GAAGGTGAAGGTCGGAGTC-3′, reverse: 5′-GAAGATGGTGATGGGATTTC-3′), *MX1* (Forward: 5′-ACAGGACCATCGGAATCTTG-3′, reverse: 5′-CCCTTCTTCAGGTGGAACAC-3′), *OAS1* (Forward: 5′-TGTCCAAGGTGGTAAAGGGTG-3′, reverse: 5′CCGGCGATTTAACTGATCCTG-3′).

### Statistical analysis

The data for figures 1–3 were analyzed using the Student’s *t*-test in Microsoft Excel. The data for figures 4, 5, and Supplemental Figure 1 were evaluated as follows: for body weight, general condition score, food intake, viral number in the lungs, NK activity, and IgA levels, the mean and standard error for each group were calculated. For food intake, the average daily food intake was calculated from the feeding day to the residual measurement day. Wilcoxon’s rank sum test was used to evaluate the general condition score. Survival analysis was performed using the log-rank test, and the analysis was adjusted to account for the multiplicity of between-group comparisons. Body weight measurements were evaluated using the Student’s *t*-test when variances were equal, and the Aspin-Welch’s *t*-test when variances were unequal, after analyzing the equivalence of variances with the F-test. Statistical analysis was performed using a commercial statistical program SAS system (SAS Institute Japan).

**Fig 1.**
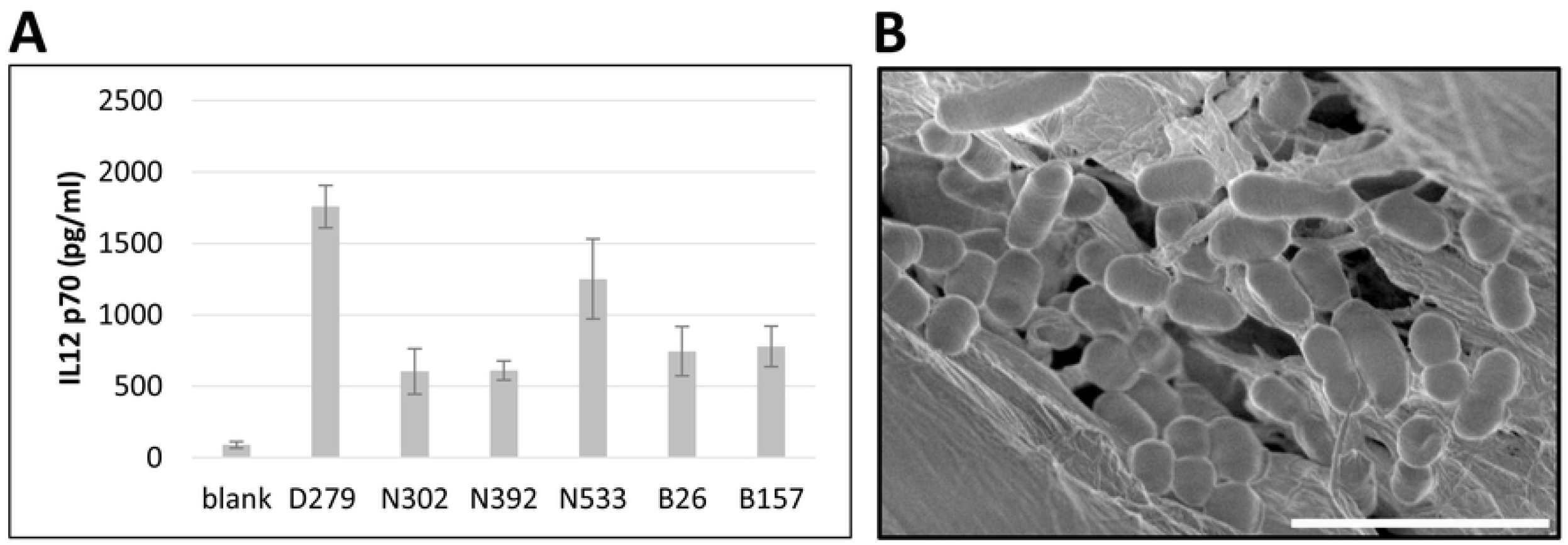
Screening the functional LAB with immunostimulatory activity. (A) IL-12 activity. Cells from the spleen of mice were cultured overnight after the introduction of LAB genomes by lipofection. The amount of IL-12 p70 produced was quantified by ELISA. The top 6 strains (>600 pg/ml) are shown in the graph. The values indicate the mean±S.E. (B) Electron microscopy image for D279. Scale bar = 5 μm.

**Fig 2.**
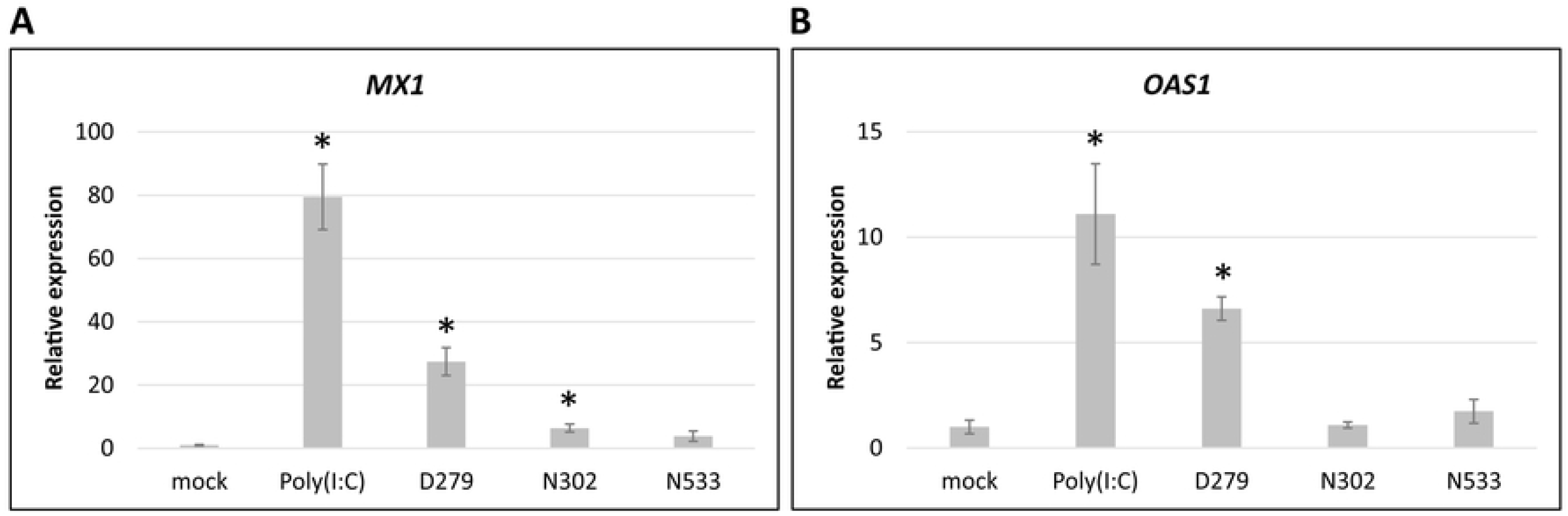
Expression analysis of anti-viral genes by quantitative RT-PCR. qRT-PCR analysis of the anti-viral genes. Twenty-four hours after introducing the LAB genomes by lipofection, total RNA was extracted from HT-29 cells and subjected to qRT-PCR. Mock (non-treated), Poly(I: C) (Poly(I: C)-treated), and LAB (LAB genome introduced) were analyzed. *Gapdh* was used as an internal control, and the results are shown relative to the expression of the mock control. Values indicate the mean±S.E. (n=4). Asterisks indicate a statistically significant difference compared with the mock control analyzed using the Student’s *t*-test (*P*<0.01). (A) Expression of *MX1*. (B) Expression of *OAS1*.

**Fig 3.**
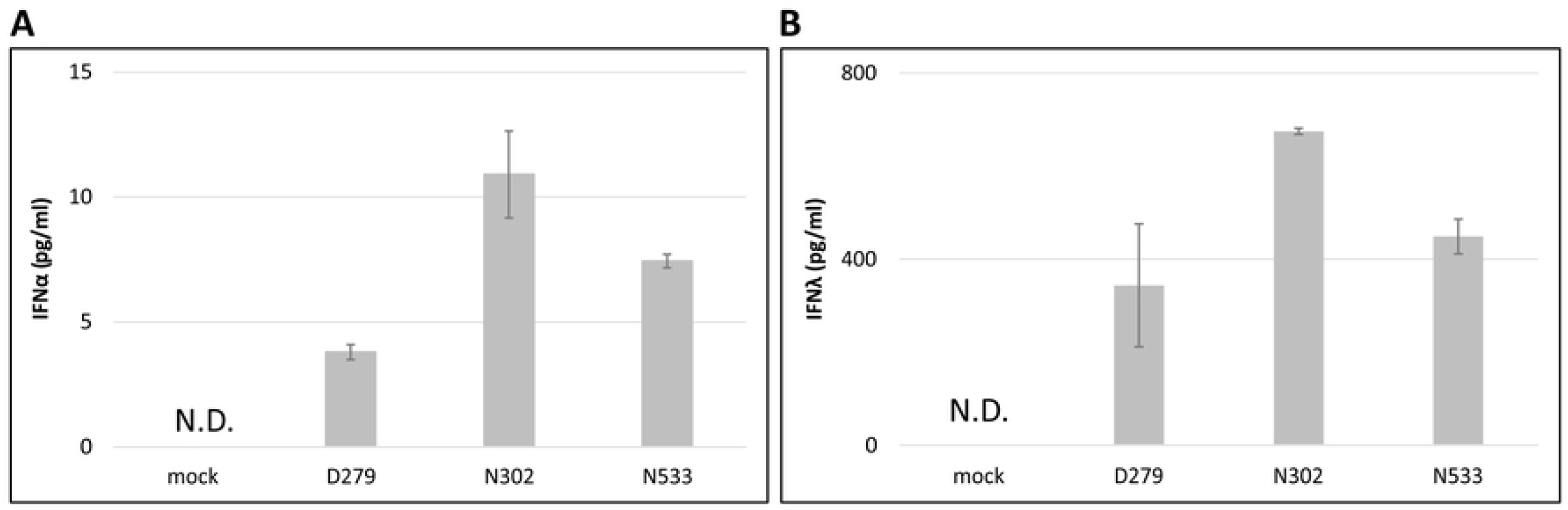
IFN induction capability of D279. LAB genomes were introduced into HT-29 cells by lipofection. After 24 h, the medium was collected and subjected to an enzyme-linked immunosorbent assay. Cultures were performed in triplicate to quantify IFNs in each experiment. Values indicate the mean±S.E. N.D. indicates not detected. (A) All IFNα subtypes. (B) All IFNλ subtypes.

**Fig 4.**
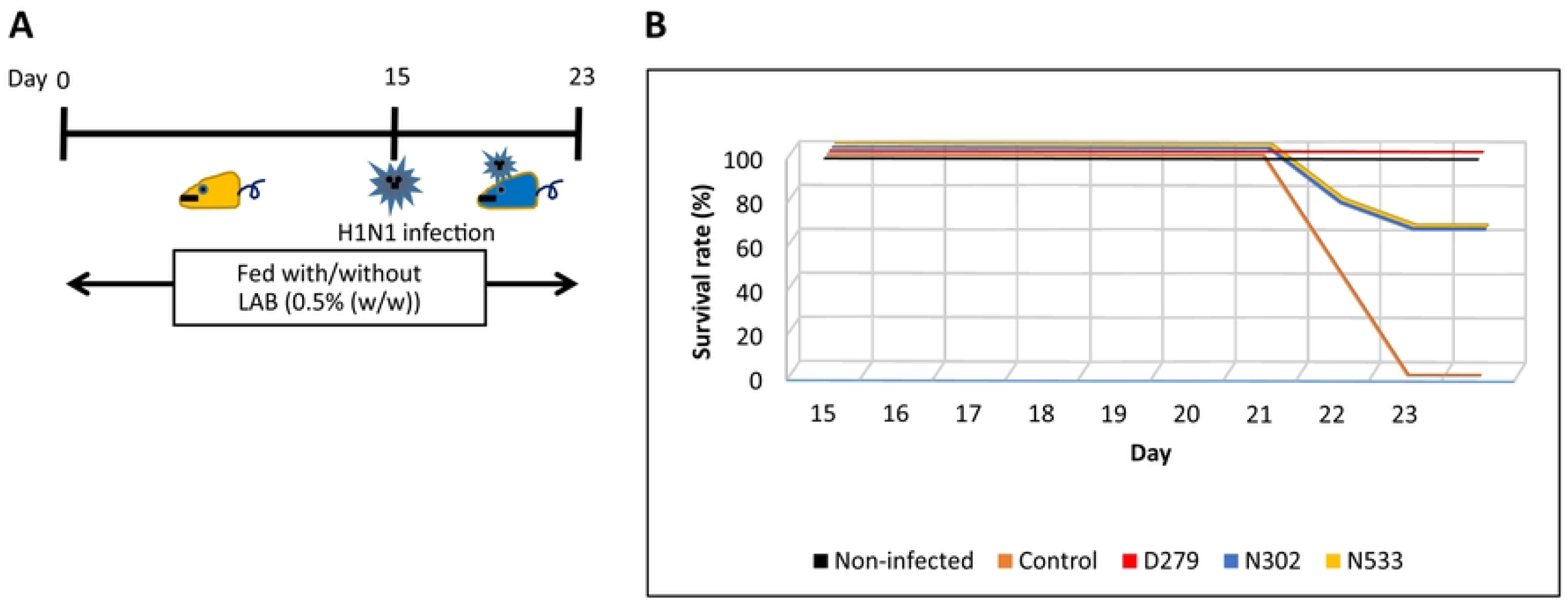
Effect of D279 during H1N1 infection testing. Experiments with a mouse influenza model were performed to investigate the immunostimulatory function of LAB. Mice were classified into non-infected, control (infected, normal feeding), and LAB-fed (infected, fed with LAB) groups (n=8 each). (A) Experimental design for the H1N1 infection test. Mice were administered food with/without LAB for 14 days before infection. On day 15, the mice were infected by dropping 50 μl of influenza virus-containing water onto the nasal cavity (1.0×10^2^ PFU/mouse). (B) Survival rates. In each group, the survival rate was monitored daily.

**Fig 5.**
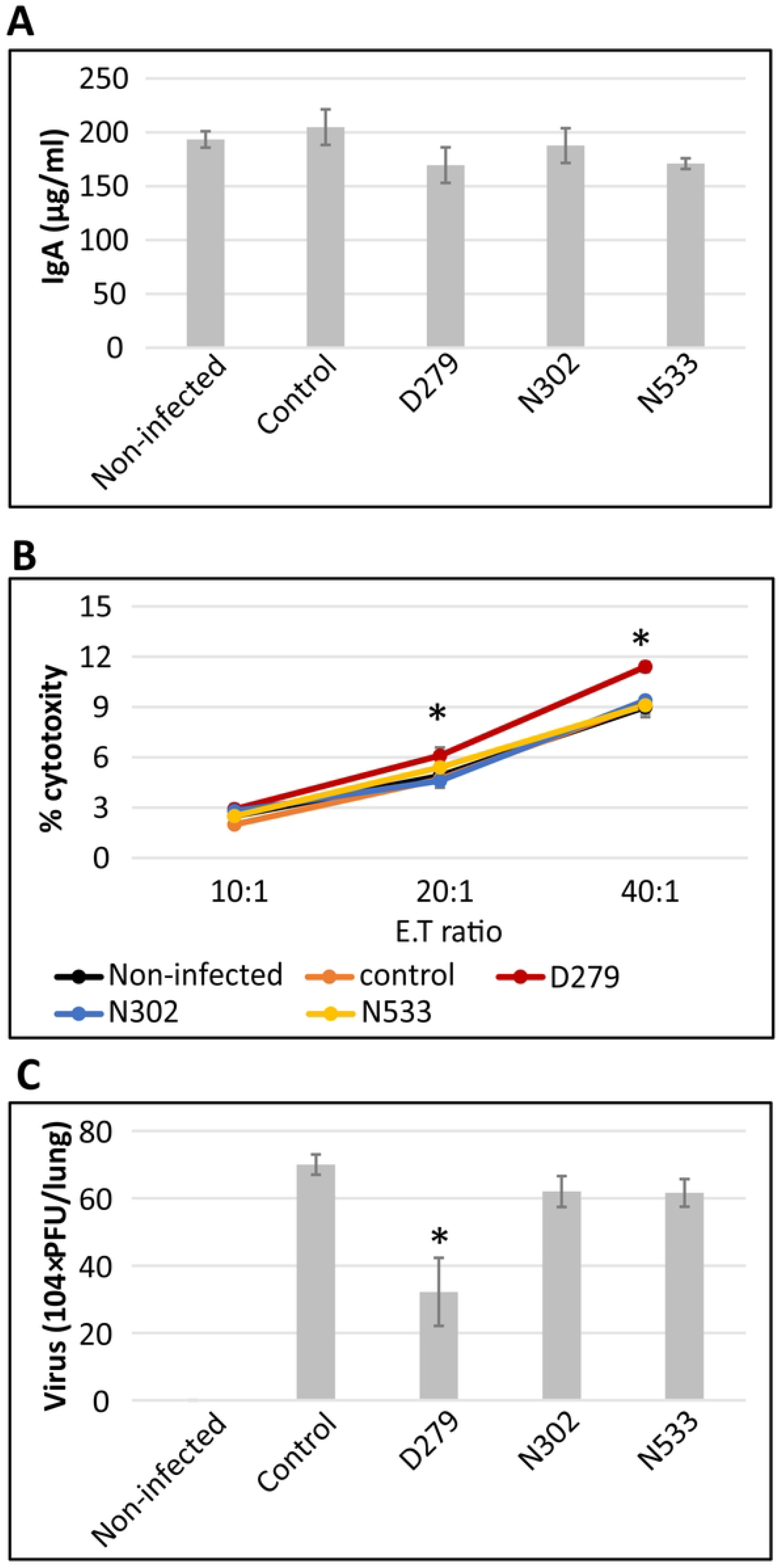
Immunostimulatory activity of D279. (A) IgA quantification. Five days after viral inoculation, serum was collected from the blood of mice and used to quantify IgA levels by ELISA. Results are shown as the mean±S.E. (n=5). (B) The number of viruses in the lung. The lungs of mice were excised after euthanasia and lung virus titers were calculated by plaque assay. The asterisk shows a significant difference using the Wilcoxon’s test (*P*<0.01) compared to the control samples. (B) NK activity. After euthanasia, mouse spleens were excised to obtain single-cell suspensions. NK activity was evaluated and the values indicate the mean±S.E. The asterisk indicates a significant difference using the Wilcoxson’s test (*P*<0.01) compared to the control. (C) Virus numbers in the lung. The number of viruses in the lung was quantified in the excised lungs. Values indicate the mean±S.E. The asterisk indicates a significant difference was obtained using the Wilcoxon’s test (*P*<0.01) compared to the control.

## Results

### Identification of LAB with improved potential to activate innate immunity

A total of 741 LAB strains were screened. The immunomodulatory ability of the LAB was evaluated based on IL-12 p70 production in splenocyte culture. IL-12 is a critical cytokine that plays a central role in activating innate immunity, and is used as a major index in LAB screening as it can activate NK cells [29, 30]. Heat-killed and lyophilized LAB were added to the splenocyte culture at a concentration of 20 μg/ml. The amount of IL-12 in the culture was quantified by ELISA. Six of the 741 strains showed significant production of IL-12 (> 600 pg/ml) (Fig 1A). Among these strains, D279 produced the highest levels of IL-12 (Fig 1A). D279 was identified as *Latilactobacillus sakei* based on phylogenetic analysis using the 16S rDNA sequence (Suppl. Fig 2), and showed a rod-shaped phenotype (Fig 1B). The D279 strain was used for subsequent experiments. For comparison, N533 (NITE_BP-03647, *Pediococcus pentosaceus*) and N302 (NITE_BP-03646, *Lactiplantibacillus plantarum*)strains were used for further analysis as N533 and N302 showed the 2^nd^ highest and lowest production of IL-12, respectively.

### Quantitative RT-PCR analysis of anti-viral genes

To obtain additional evidence for the immunostimulatory function of D279, the expression levels of the anti-viral genes *MX1* (MX dynamin-like GTPase1) and *OAS1* (2′, 5′-oligoadenylate synthase1) were analyzed *in vitro*, as these genes inhibit viral gene replication [34, 35]. HT-29 cells were cultured overnight in 96-well plates and the genomes of D279, N302, and N533 were introduced by lipofection. Total RNA was extracted after 24 h of culture, and analyzed by qRT-PCR. The expression of *MX1* and *OAS1* was highly induced in genomic DNA-introduced cells; however, the D279 genome induced the highest expression of these genes among the LAB analyzed (Fig 2A, 2B). The result indicates that D279 showed strong immunostimulatory activity *in vitro*.

### IFN production in vitro

To investigate the mode of action of D279, Type I and Type III IFN production were analyzed *in vitro* because of their pivotal role in viral defense. HT-29 cells were grown in a 96-well plate for 24 h, and the LAB genomes were introduced by lipofection. N302 and N533 significantly induced IFNα and IFNλ compared to the mock control (Fig 3A, 3B). In HT-29 cells transfected with the D279 genome, IFNα was produced, but the levels were significantly lower than that of N302 and N533 (Fig 3A), whereas IFNλ was induced at similar levels as N533. The ratio of IFNα to IFNλ was 1:90 in D279, whereas that of N302 and N533 was 1:60, indicating that D279 preferentially induces Type III IFN. Thus, D279 stimulated innate immunity in a less inflammatory manner than the other LAB strains analyzed.

### Influenza infection test

To test the biogenic effects of D279, a mouse influenza (H1N1) infection model was used. Mice were fed *ad libitum* with or without LAB powder mixed at 0.5% (w/w) for 14 days before influenza infection (Fig 4A). The mice were divided equally into three groups (n=8 per group): non-infected (no infection, normal feeding), control (infected, normal feeding), and LAB-fed (infected, fed with D279, N302, and N533). On day 15, the mice were infected with the influenza virus by dropping the influenza-containing medium onto the nasal cavity (1.0×10^2^ PFU/mouse) (Fig 4A). The test was continued until day 23 (Fig 4A). In the control group, half the mice were euthanized on day 21, and the other half were euthanized on day 22 according to humanitarian endpoints, whereas all mice survived in the non-infected group (Fig 4B). N302 and N533 showed anti-viral function, as the survival rate improved to 63% on day 23. Importantly, the D279 administered group showed a 100% survival rate during the test period (Fig 3B), indicating greater beneficial effects than the other LAB strains.

Additionally, mouse phenotypes were observed daily and evaluated for clinical scores (Suppl. Fig 1A, Table 1). The score increased when the symptoms worsened after infection (Table 1). The D279 administered group showed a slower increase in the clinical score than the other groups (Suppl. Fig 1A), whereas body weight and food intake continued to decrease throughout the study period (Suppl. Fig 1B, 2C), suggesting that the higher survival rate of the D279-fed group was related to the immunostimulatory activity of D279. These results indicate that oral administration of D279 before influenza infection is effective in preventing viral invasion.

### The mechanism of immunostimulation by D279

*In vivo* experiments demonstrated that D279 alleviated influenza symptoms. To determine the reason for the higher survival rate in the D279-fed group, the degree of activation of humoral and cellular immunity was examined. The amount of IgA was quantified; however, its production did not differ among the groups (Fig 5A), indicating that D279 did not stimulate humoral immunity. Next, NK activity was quantified by evaluating the number of NK cells that attacked FITC-labeled cancer cells *in vitro*. The LAB-fed groups showed higher NK activity; however, in the D279-fed group, NK activity was significantly higher than that in the other LAB-fed groups (Fig 5B). Further, the number of viruses in the lungs was quantified using a plaque assay. The D279-fed group alone showed a significant reduction in viral number (Fig 5C). These data indicate that D279 stimulates cellular, but not humoral immunity *in vivo*.

## Discussion

In this study, D279 was identified as a useful strain that conferred a 100% survival rate in influenza infection tests in mice. D279 was isolated from among 741 strains of IL-12-producing LAB. D279 was identified as *L. sakei*, and its immunostimulatory activity has not been evaluated. Several strains of the genus *L. sakei* have been isolated from processed meat, fish, and vegetable products, and are traditionally used as fermentation starters [36, 37]. The type strain of *L. sakei* ATCC15521 (subsp. *sakei*) was isolated more than 80 years ago, and has been shown to have immunostimulatory functions [38]. D279 is a plant lactobacillus that was originally isolated from the Japanese pickles Aka-kabu zuke (a red turnip pickled with LAB and with/without salts). The pickle is a traditional food that has long been consumed during winter when influenza and other respiratory viruses are present. It is interesting to note the traditional methods that people have adopted in their daily diet to prevent viral infection. In addition, some LAB of the genus *L. sakei* reportedly activate mucosal immunity [39–41]. Thus, *L. sakei* appears to be highly beneficial for human health.

The oral administration of D279 to mice dramatically improved their survival rate after influenza infection, and no mice showed mortality during the experimental period, whereas the other LAB-fed groups showed a reduced survival rate. The improvement in survival rates resulted due to the immunostimulatory effect of D279, as dietary intake did not change between groups administered D279 or not (Suppl. Fig 1C). Several LAB have anti-viral effects depending on the strains used [42, 43]. For example, oral administration of *Lactobacillus pentosus* ONRICb240, *Lactobacillus delbrueckii* OLL1073R-1, *Lactobacillus casei* strain Shirota, *Lactobacillus crispatus* KT-11, and *Lactobacillus pentosus* strain S-PT84 to mice resulted in anti-viral effects due to the activation of innate immunity via the production of macrophages, NK cells, IgA, Type I IFN, TNFα, and IL-6 after Th1 polarization [44–48]. The macrophage-NK cell pathway is frequently activated by LAB and is a key barrier against viral infection. *Lactococcus lactis* JCM5805 activates innate immunity by directly stimulating pDCs to promote type I IFN production [19]. D279 administration increased NK cell activity, but not IgA production, during the influenza infection test (Fig 5A, 5C), and D279 may show uptake by macrophages, which may in turn stimulate IL-12 production and activation of NK cells. IgA production requires Type I IFN from pDCs, B cell activating factor from the TNF family (BAFF), and a proliferation inducing ligand (APRIL) to stimulate B cells [48, 49, 50]. Because D279 does not significantly induce IgA production, uptake by pDCs and their activation may not occur. Thus, oral administration of D279 directly eliminated influenza viruses by driving cellular immunity, including macrophages and NK cells.

Importantly, D279 stimulates IFN production. The ratio of IFNα:IFNλ was 1:90 in D279, whereas that produced by N302 and N533 was 1:60. The result indicates that D279 preferentially induces IFNλ to activate innate immunity, resulting in a relatively low inflammatory response. IFNα and IFNλ induce the expression of the same subset of anti-viral genes; however, the distinct role of IFNλ has been identified recently. Davidson et al. demonstrated the effectiveness of IFNλ, rather than IFNα, against respiratory viruses, and their results showed that administration of IFNα to influenza-infected cells or mice reduced the viral load, although inflammatory stress resulted in necrosis or death [17]. However, IFNλ administration effectively decreased the number of virus particles, and a better survival rate was observed than with IFNα treatment. Dramatic improvement in the survival of the D279-fed group may result in part due to reduced inflammatory immune activation. However, one limitation of this study is that we did not obtain enough data on cytokine production, including IL-12, IFNα, and IFNλ *in vivo*, because of the limited number of mice for in-depth analysis. Further *in vivo* studies are required to quantify IFN production, and that of other inflammatory and noninflammatory cytokines.

In future studies, it will be important to elucidate the immunostimulatory mechanism of D279. The components of D279 that are activated and how they affect innate immunity to result in a 100% survival rate are unclear. Thus, D279 exhibited anti-viral effects, making it a valuable strain for commercial use. Importantly, D279 can be used as a food additive or supplement for health maintenance.

## Supporting information

Supplementary Fig 1: Clinical outcomes.

Supplementary Fig 2: Phylogenetic analysis.

## Acknowledgments

We would like to thank Japan Bioresearch Center Co., Ltd. for performing and analyzing the animal tests. We also thank TechnoSuruga Laboratory Co., Ltd. for analyzing the 16S rDNA sequences. We would like to thank Editage (www.editage.com) for English language editing.

## Supplementary Figures

**Supplementary Fig 1: Classification of D279**

Maximum-likelihood phylogenetic tree of D279 with related strains. The 16S rDNA sequence was aligned, and bootstrap analysis was performed with 1000 replicates. The scale bar indicates 0.01 substitutions per nucleotide position.

**Supplemental Fig 2: Clinical outcomes**

For the influenza model experiments, the clinical scores, survival rates, feeding amount, and body weight were measured every day until day 23. (A) Clinical scores. From day 0 to day 23, phenotypes were observed and scored according to the severity of the body condition. A higher score indicates a more severe condition. The mice in the control group were sacrificed on day 22 as per the humanitarian endpoint. (B) Body weight. During the H1N1 infection experiment, the body weights of the surviving mice were measured daily. The values shown indicate the mean±S.E. (C) Food consumption. Changes in food consumption were measured twice a week. On the day of feeding, the amount of food in each feeding container was weighed with an electronic balance, and the remaining amount was measured on the next feeding day. Values indicate the mean±S.E.Figure

## References

1. Perdigón G, Fuller R, Raya R. Lactic acid bacteria and their effect on the immune system. Curr Issues Intest Microbiol. 2001;2: 27–42.

2. Ren C, Faas, MM, de Vos P. Disease managing capacities and mechanisms of host effects of lactic acid bacteria. Crit Rev Food Sci Nutr. 2021;61: 1365–1393. doi: 10.1080/10408398.2020.1758625.

3. Ayivi RD, Gyawali R, Krastanov A, Aljaloud S.O, Worku M, Tahergorabi R, et al. Lactic Acid Bacteria: Food safety and human health applications. Dairy. 2020;1: 202–232. doi: 10.3390/dairy1030015.

4. Kanauchi O, Andoh A, AbuBakar S, Yamamoto N. Probiotics and paraprobiotics in viral infection: clinical application and effects on the innate and acquired immune systems. Curr Pharm Des. 2018;24: 710–717. doi: 10.2174/1381612824666180116163411.

5. Watanabe T, Hayashi K, Takahashi I, Ohwaki M, Kan T, Kawahara T. Physical properties of lactic acid bacteria influence the level of protection against influenza infection in mice. PLoS One. 2021;16: e0251784. doi: 10.1371/journal.pone.0251784

6. Quinto EJ, Jiménez P, Caro I, Tejero J, Mateo J, Girbés T. Probiotic lactic acid bacteria: A review. Food Nutr Sci. 2014;5: 1765–1775. doi: 10.4236/fns.2014.518190.

7. Perdigón G, Vintiñi E, Alvarez S, Medina M, Medici M. Study of the possible mechanisms involved in the mucosal immune system activation by lactic acid bacteria. J Dairy Sci. 1999;82: 1108–1114. doi: 10.3168/jds.S0022-0302(99)75333-6.

8. Granucci F, Foti M, Ricciardi-Castagnoli P. Dendritic cell biology. Adv Immunol. 2005;88: 193–233. doi: 10.1016/S0065-2776(05)88006-X.

9. Fu YL, Harrison RE. Microbial phagocytic receptors and their potential involvement in cytokine induction in macrophages. Front Immunol. 2021;12: 662063. doi: 10.3389/fimmu.2021.662063.

10. Stuart LM, Paquette N, Boyer L. Effector-triggered versus pattern-triggered immunity: How animals sense pathogens. Nat Rev Immunol. 2013;13: 199–206. doi: 10.1038/nri3398.

11. Liu Q, Ding JL. The molecular mechanisms of TLR-signaling cooperation in cytokine regulation. Immunol Cell Biol. 2016;94: 538–542. doi: 10.1038/icb.2016.18.

12. Kawai T, Akira S. TLR signaling. Semin Immunol. 2007;19: 24–32. doi: 10.1016/j.smim.2006.12.004.

13. Vidal SM, Khakoo SI, Biron CA. Natural killer cell responses during viral infections: Flexibility and conditioning of innate immunity by experience. Curr Opin Virol. 2011;1: 497–512. doi: 10.1016/j.coviro.2011.10.017.

14. Chen K, Liu J, Cao Xuetao. Regulation of type I interferon signaling in immunity and inflammation: A comprehensive review. J Autoimmun. 2017;83: 1–11. doi: 10.1016/j.jaut.2017.03.008.

15. Syedbasha M, Egli A. Interferon lambda: Modulating immunity in infectious diseases. Front Immunol. 2017;8: 119. doi: 10.3389/fimmu.2017.00119.

16. McNab F, Mayer-Barber K, Sher A, Wack A, O’Garra A. Type I interferons in infectious disease. Nat Rev Immunol. 2015;15: 87–103. doi: 10.1038/nri3787.

17. Davidson S, McCabe TM, Crotta S, Gad HH, Hessel EM, Beinke S, et al. IFNλ is a potent anti-influenza therapeutic without the inflammatory side effects of IFNα treatment. EMBO Mol Med. 2016;8: 1099–1112. doi: 10.15252/emmm.201606413.

18. Lasfar A, Zloza A, de la Torre A, Cohen-Solal KA. IFN-λ: A new inducer of local immunity against cancer and infections. Front Immunol. 2016;7: 598. doi: 10.3389/fimmu.2016.00598.

19. Jounai K, Ikado K, Sugimura T, Ano Y, Braun J, Fujiwara D. Spherical lactic acid bacteria activate plasmacytoid dendritic cells immunomodulatory function via TLR9-dependent crosstalk with myeloid dendritic cells. PLoS One. 2012;7: e32588. doi: 10.1371/journal.pone.0032588.

20. Sugimura T, Jounai K, Ohshio K, Tanaka T, Suwa M, Fujiwara D. Immunomodulatory effect of Lactococcus lactis JCM5805 on human plasmacytoid dendritic cells. Clin Immunol. 2013;149: 509–518. doi: 10.1016/j.clim.2013.10.007.

21. Suzuki H, Jounai K, Ohshio K, Fujii T, Fujiwara D. Administration of plasmacytoid dendritic cell-stimulative lactic acid bacteria enhances antigen-specific immune responses. Biochem Biophys Res Commun. 2018;503: 1315–1321. doi: 10.1016/j.bbrc.2018.07.042.

22. Iliev ID, Kitazawa H, Shimosato T, Katoh S, Morita H, He F, et al. Strong immunostimulation in murine immune cells by Lactobacillus rhamnosus GG DNA containing novel oligodeoxynucleotide pattern. Cell Microbiol. 2005;7: 403–414. doi: 10.1111/j.1462-5822.2004.00470.x.

23. Inoue R, Nagino T, Hoshino G, Ushida K. Nucleic acids of Enterococcus faecalis strain EC-12 are potent toll-like receptor 7 and 9 ligands inducing interleukin-12 production from murine splenocytes and murine macrophage cell line J774.1. FEMS Immunol Med Microbiol. 2011;61: 94–102. doi: 10.1111/j.1574-695X.2010.00752.x.

24. Gao K, Wang C, Liu L, Dou X, Liu J, Yuan L, et al. Immunomodulation and signaling mechanism of Lactobacillus rhamnosus GG and its components on porcine intestinal epithelial cells stimulated by lipopolysaccharide. J Microbiol Immunol Infect. 2017;50: 700–713. doi: 10.1016/j.jmii.2015.05.002.

25. Marongiu L, Gornati L, Artuso I, Zanoni I, Granucci F. Below the surface: The inner lives of TLR4 and TLR9. J Leukocyte Biol. 2019;106: 147–160. doi: 10.1002/JLB.3MIR1218-483RR.

26. Miettinen M, Veckman V, Latvala S, Sareneva T, Matikainen S, Julkunen I. Live lactobacillus rhamnosus and streptococcus pyogenes differentially regulate Toll-like receptor (TLR) gene expression in human primary macrophages. J Leukocyte Biol. 2008;84: 1092–1100. doi: 10.1189/jlb.1206737.

27. Ramani T, Auletta VS, Weinstock D, Mounho-Zamora B, Ryan PC, Salcedo TW, et al. Cytokines: The Good, the Bad, and the Deadly. Int J Toxicol. 2015;34: 355–365 doi: 10.1177/1091581815584918.

28. Gu Y, Zuo X, Zhang S, Ouyang Z, Jiang S, Wang F, et al. The mechanism behind influenza virus cytokine storm. Viruses. 2021;13: 1362–1378. doi: 10.3390/v13071362.

29. Guo Y, Cao W, Zhu Y. Immunoregulatory functions of the IL-12 family of cytokines in antiviral systems. Viruses. 2019;11: 772–784. doi: 10.3390/v11090772.

30. Monteiro JM, Harvey C, Trinchieri G. Role of interleukin-12 in primary influenza virus infection. J Virol. 1998;72: 4825–4831. doi: 10.1128/JVI.72.6.4825-4831.1998.

31. Hama Y, Kurokawa M, Imakita M, Yoshida Y, Shimizu T, Watanabe W, et al. Interleukin 12 is a primary cytokine responding to influenza virus in the respiratory tract of mice. Acta Virol. 2009;53: 233–240. doi: 10.4149/av_2009_04_233.

32. Chang HD, Radbruch A. The pro- and anti-inflammatory potential of interleukin-12. Ann N Y Acad Sci. 2007; 1109:40–6, doi: 10.1196/annals.1398.006.

33. Thangavel RR, Bouvier NM. Animal models for influenza virus pathogenesis, transmission, and immunology. J Immunol Methods. 2014;410: 60–79. doi: 10.1016/j.jim.2014.03.023.

34. Verhelst J, Parthoens E, Schepens B, Fiers W, Saelens X. Interferon-inducible protein Mx1 inhibits influenza virus by interfering with functional viral ribonucleoprotein complex assembly. J Virol. 2012;86: 13445–13455. doi: 10.1128/JVI.01682-12.

35. Kristiansen H, Scherer CA, McVean M, Iadonato SP, Vends S, Thavachelvam K, et al. Extracellular 2’-5’ oligoadenylate synthetase stimulates RNase L-independent antiviral activity: A novel mechanism of virus-induced innate immunity. J Virol. 2010;84: 11898–11904. doi: 10.1128/JVI.01003-10.

36. Ammor S, Dufour E, Zagorec M, Chaillou S, Chevallier I. Characterization and selection of Lactobacillus sakei strains isolated from traditional dry sausage for their potential use as starter cultures. Food Microbiol. 2005;22: 529–538. doi: 10.1016/j.fm.2004.11.016.

37. Zagorec M, Champomier-Vergès M-C. Lactobacillus sakei: A Starter for Sausage Fermentation, a Protective Culture for Meat Products. Microorganisms. 2017;5: 56. doi: 10.3390/microorganisms5030056.

38. Katagiri H, Kitahara K, Fukami K. The characteristics of the lactic acid bacteria isolated from Moto, yeast mashes for saké manufacture. Bull Agric Chem Soc Jpn. 1934;10: 153–157.

39. Mathiesen G, Huehne K, Kroeckel L, Axelsson L, Eijsink VGH. Characterization of a new bacteriocin operon in sakacin P-producing Lactobacillus sakei, showing strong translational coupling between the bacteriocin and immunity genes. Appl Environ Microbiol. 2005;71: 3565–3574. doi: 10.1128/AEM.71.7.3565-3574.2005.

40. Yamasaki-Yashiki S, Miyoshi Y, Nakayama T, Kunisawa J, Katakura Y. IgA-enhancing effects of membrane vesicles derived from Lactobacillus sakei subsp. sakei NBRC15893. Bioscience of Microbiota. Biosci Microbiota Food Health. 2019;38: 23–29. doi: 10.12938/bmfh.18-015.

41. Ghoneum M, Abdulmalek S. KDP, a lactobacilli product from kimchi, enhances mucosal immunity by increasing secretory IgA in mice and exhibits antimicrobial activity. Nutrients. 2021;13: 3936. doi: 10.3390/nu13113936.

42. Nagafuchi S, Takahashi T, Yajima T, Kuwata T, Hirayama K, Itoh K. Strain dependency of the immunopotentiating activity of Lactobacillus delbrueckii subsp. bulgaricus. Biosci Biotechnol Biochem. 1999;63: 474–479. doi: 10.1271/bbb.63.474.

43. Fujiwara D, Inoue S, Wakabayashi H, Fujii T. The anti-allergic effects of lactic acid bacteria are strain dependent and mediated by effects on both Th1/Th2 cytokine expression and balance. Int Arch Allergy Immunol. 2004;135: 205–215. doi: 10.1159/000081305.

44. Kotani Y, Kunisawa J, Suzuki Y, Sato I, Saito T, Toba M, et al. Role of Lactobacillus pentosus Strain b240 and the toll-like receptor 2 axis in Peyer’s patch dendritic cell-mediated immunoglobulin A enhancement. PLOS One. 2014;9: e91857. doi: 10.1371/journal.pone.0091857.

45. Makino S, Ikegami S, Kano H, Sashihara T, Sugano H, Horiuchi H, et al. Immunomodulatory effects of polysaccharides produced by Lactobacillus delbrueckii ssp. bulgaricus OLL1073R-1. J Dairy Sci. 2006;89: 2873–2881. doi: 10.3168/jds.S0022-0302(06)72560-7.

46. Yasui H, Kiyoshima J, Hori T. Reduction of influenza virus titer and protection against influenza virus infection in infant mice fed Lactobacillus casei Shirota. Clin Diagn Lab Immunol. 2004;11: 675–679. doi: 10.1128/CDLI.11.4.675-679.2004.

47. Tobita K, Yanaka H, Otani H. Lactobacillus crispatus KT-11 enhances intestinal immune functions in C3H/HeN mice. J Nutr Sci Vitaminol (Tokyo). 2010;56: 441–445. doi: 10.3177/jnsv.56.441.

48. Koizumi S, Wakita D, Sato T, Mitamura R, Izumo T, Shibata H, et al. Essential role of Toll-like receptors for dendritic cell and NK1.1(+) cell-dependent activation of type 1 immunity by Lactobacillus pentosus strain S-PT84. Immunol Lett. 2008;120: 14–19. doi: 10.1016/j.imlet.2008.06.003.

49. Litinskiy MB, Nardelli B, Hilbert DM, He B, Schaffer A, Casali P, et al. DCs induce CD40-independent immunoglobulin class switching through BLyS and APRIL. Nat Immunol. 2002;3: 822–829. doi: 10.1038/ni829.

50. He B, Xu W, Santini PA, Polydorides AD, Chiu A, Estrella J, et al. Intestinal bacteria trigger T cell-independent immunoglobulin A(2) class switching by inducing epithelial-cell secretion of the cytokine APRIL. Immunity. 2007;26: 812–826. doi: 10.1016/j.immuni.2007.04.014.

